# Comparative Qualitative Phosphoproteomics Analysis Identifies Shared Phosphorylation Motifs and Associated Biological Processes in Flowering Plants

**DOI:** 10.1101/233668

**Authors:** Shireen Al-Momani, Da Qi, Zhe Ren, Andrew R Jones

## Abstract

Phosphorylation is regarded as one of the most prevalent post-translational modifications and plays a key role in regulating cellular processes. In this work we carried out a comparative bioinformatics analysis of phosphoproteomics data, to profile two model species representing the largest subclasses in flowering plants the dicot *Arabidopsis thaliana* and the monocot *Oryza sativa*, to understand the extent to which phosphorylation signaling and function is conserved across evolutionary divergent plants. Using pre-existing mass spectrometry phosphoproteomics datasets and bioinformatic tools and resources, we identified 6,537 phosphopeptides from 3,189 phosphoproteins in *Arabidopsis* and 2,307 phosphopeptides from 1,613 phosphoproteins in rice. The relative abundance ratio of serine, threonine, and tyrosine phosphorylation sites in rice and *Arabidopsis* were highly similar: 88.3: 11.4: 0.4 and 86.7: 12.8: 0.5, respectively. Tyrosine phosphorylation shows features different from serine and threonine phosphorylation and was found to be more frequent in doubly-phosphorylated peptides in *Arabidopsis*. We identified phosphorylation sequence motifs in the two species to explore the similarities, finding nineteen pS motifs and two pT motifs that are shared in rice and *Arabidopsis*; among them are five novel motifs that have not previously been described in both species. The majority of shared motif-containing proteins were mapped to the same biological processes with similar patterns of fold enrichment, indicating high functional conservation. We also identified shared patterns of crosstalk between phosphoserines with motifs pSXpS, pSXXpS and pSXXXpS, where X is any amino acid, in both species indicating this is an evolutionary conserved signaling mechanism in flowering plants. However, our results are suggestive that there is greater co-occurrence of crosstalk between phosphorylation sites in *Arabidopsis*, and we were able to identify several pairs of motifs that are statistically significantly enriched to co-occur in *Arabidopsis* proteins, but not in rice.

## Introduction

Phosphorylation is regarded as one of the most prevalent post-translational modifications (1). The reaction is catalyzed by a protein kinase to transfer the γ-phosphoryl group from adenosine triphosphate (ATP) or guanosine triphosphate (GTP), most commonly via a covalent bond to the hydroxyl group of a specific serine, threonine, or tyrosine amino acid within the target protein (2). One feature of protein phosphorylation that makes it an ideal participant in signal transduction pathways is the reversibility of its chemical reaction through the subsequent removal of the phosphoryl group attached to the phosphorylated amino acid by a protein phosphatase, allowing for signal transduction cascade to maintain a prompt cellular response to stimuli coming from outside or within the cells (3).

Phosphorylation regulates protein function and cell signaling by triggering a change in the three-dimensional structure of the protein, which in turn influences the protein into behaving differently by activating or deactivating its catalytic function. A change in the structure of a phosphorylated protein (pProtein) can also recruit other proteins that contain structurally conserved domains to recognize and bind to specific motifs (4, 5). Phosphorylation events are crucial to understanding the functional biology of plants, since they control essential biological processes including seed germination, stomatal movement, pistil development and pollination, the innate immune response, defense and stress tolerance (6, 7, 8, 9, 10).

Monocots and dicots are the largest subclasses in flowering plants (Angiosperms) (11). The monocot lineage branched off from dicots approximately 140-150 million years ago (12), yet many key mechanisms and transcription factors present in both dicots and monocots regulate the expression of biotic and abiotic stress response genes (13, 14, 15, 16, 17, 18). In this work, we explore the extent to which phosphorylation-mediated signaling events are conserved, or have diverged, between two key model species – rice and *Arabidopsis*. We thus aim to understand the extent to which findings about signaling in these model organisms are likely to be transferable to other non-model plants. A past study by Nakagami et al. (2010) identified some conserved phosphorylation sites in plants based on analyzing orthologous phosphoproteins in rice and *Arabidopsis* (19). In their analysis, around 50% of pProteins in either species had an ortholog that was also phosphorylated (50.4% rice pProteins to *Arabidopsis* orthologue; 56.2% *Arabidopsis* pProtein to rice orthologue), and around 20% phosphorylated at the same site (18% rice phospho-site,· 25% *Arabidopsis* phospho-site). Our work extends from this study by examining the extent to which phosphorylation motifs are conserved in the two species, the functional associations of those motifs, and the evidence for crosstalk between proximal phospho-sites.

Rice (*Oryza sativa*) is a primary monocot model plant for cereal research due to its compact genome and evolutionary relationships with other cereals that have larger genomes (20). According to the Ensembl plants website (Assembly: IRGSP-1.0, INSDC Assembly GCA_001433935.1) *O. sativa* Japonica has a 374 MB genome, 35,679 coding genes and 48,950 protein sequences recorded in UniProt (March 2017, UniProt Proteome ID: P000059680). *O. sativa* Indica has a 412 MB genome, 40,745 coding genes in Ensembl (database version 88.2, ASM465v1, INSDC Assembly GCA_000004655.2, Jan 2005) with 37,385 protein sequences in UniProt (UP000007015). *Arabidopsis thaliana* has a 136 MB genome, 27,655 coding genes recorded in Ensembl plants (Database version 88.11, TAIR10, INSDC Assembly GCA_000001735.1, Sep 2010,) and 32,113 protein sequences in UniProt (Proteome ID UP000006548). Discrepancies between counts of coding genes and protein sequences are likely due to different genome annotations present in different resources, however, for the purposes of comparison we can conclude that the rice genome (both sub-species) is around three times larger than *A. thaliana*, and likely has 30-50% higher gene count. At present, there are 9 wild rice varieties (other *Oryza* species) with genomes in Ensembl, with gene counts ranging from ∼29K (*Oryza meridionalis*) up to 37K (*Oryza rufipogon*), indicating that the higher gene count is relatively stable and not a result of recent domestication as in the case of the wheat genome (104K genes).

The rice and *Arabidopsis* genomes contain 1,512 (in Japonica; 1403 in Indica) and 1,052 protein kinases, respectively, more than twice the number in humans (516) (21). Unique protein kinases that are only found in higher plants, such as RLKs, are positive regulators for plant tolerance to salt and cold stresses (22, 23, 24). However, not all duplicates have been retained: for example, plant protein kinases that are involved in housekeeping functions (such as metabolism, mitosis, and other primary functions that all living organisms need to survive) are kept in low copy numbers while other plant protein kinases, in particular kinases that are expressed only at specific developmental stages, have a high copy number of duplicates. It has previously been suggested that many of these protein kinases were duplicated and retained owing to their roles in plant-specific processes, while housekeeping genes might be under strong purifying selection acts to preserve their normal function (25).

In terms of experimental data on plant species, *Arabidopsis* has the most extensive proteomic data available in plants. It is one of the top ten species in the number of submitted data sets in ProteomeXchange consortium (26). Compared with *Arabidopsis*, fewer data are available in other plant species, and thus it is important to understand the extent to which findings in this model organism can be transferred to other plant species, in particular crops that are important for food security, such as rice.

In this study, we carried out a comparative qualitative phosphoproteomics analysis in dicot *A. thaliana* and monocot *O. sativa*, using pre-existing mass spectrometry (MS) phosphoproteomics datasets and several bioinformatic tools and resources. The aims of our study were to identify and compare features of rice and *Arabidopsis* phosphoproteomes, and to identify shared phosphorylation motifs and associated biological processes across flowering plants. This could enhance our understanding of regulatory systems, which in the longer term could potentially lead to improving the productivity of agronomically important plant species.

## Experimental Procedures

### Datasets

Two rice *O. sativa* Indica (PXD002222 and PXD000923) and two mouse-ear cress *A. thaliana* (PXD000033 and PXD000421) data sets were chosen from experiments that include phospho-enrichment steps (27, 28, 29, 30). MGF files for each data set were downloaded from the ProteomeXchange central website (http://www.proteomexchange.org). If MGF files were not available, raw files were converted to MGF using MSConvertGUI (31).

### Proteins and Peptides Identification and Modification Localization

PEAKS software version 7.5 (32) was used to search the spectra obtained from datasets PXD000033 and PXD000421 against the *A. thaliana* protein database and spectra obtained from datasets PXD002222 and PXD000923 against *O. sativa* Indica database. Databases were downloaded from UniProt version 59 including canonical and isoform protein sequences (33). Trypsin was specified as the proteolytic enzyme of choice, with one non-specific cleavage and three maximum variable modifications per peptide for all data sets. Other search parameters were selected to match as closely as possible the parameters from the original analysis - a summary of search parameters and cutoff filter used in PEAKS database searching is shown in Table 1.

**Table 1.**
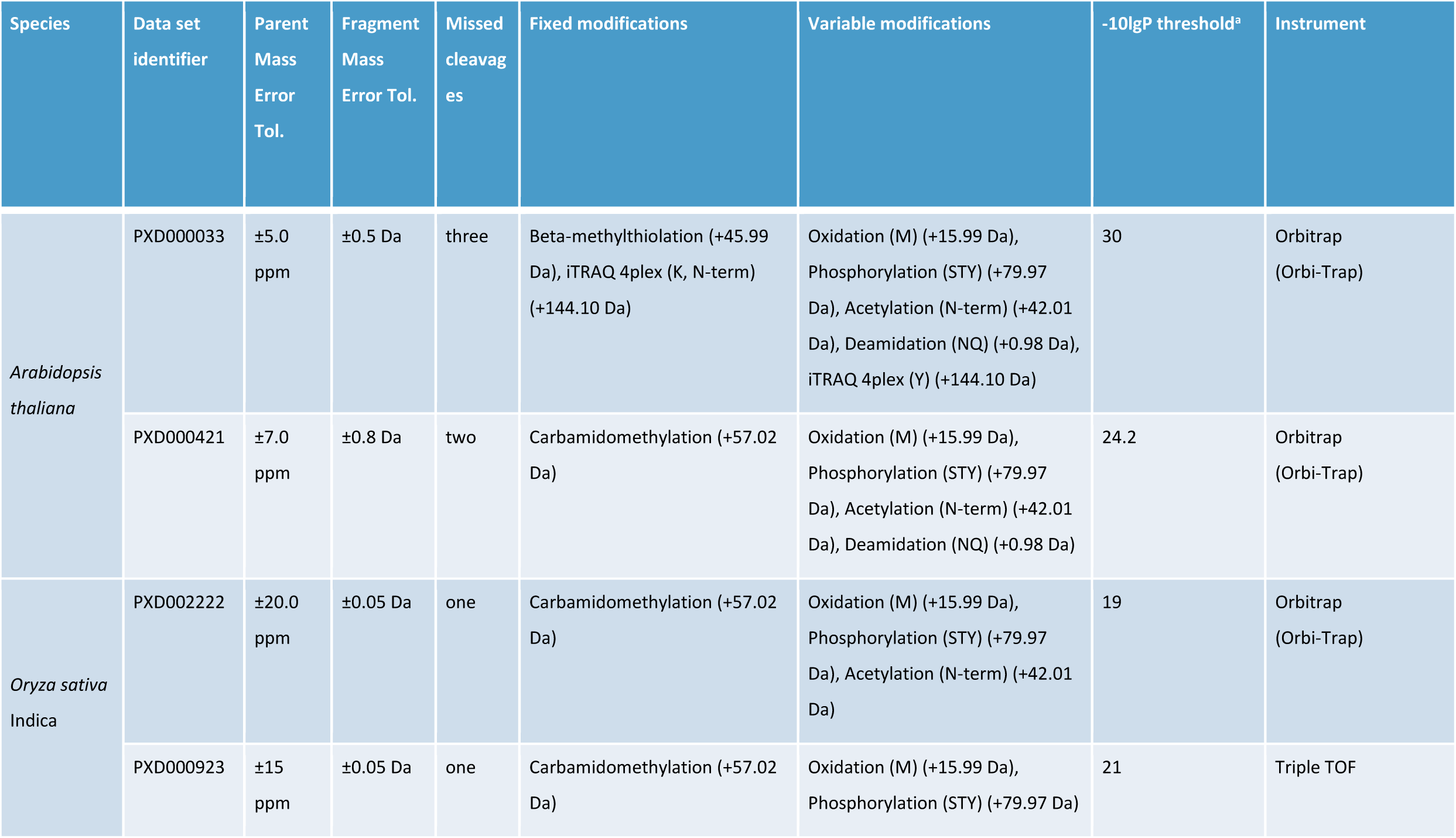
Search parameters and cutoff filter used in PEAKS database searching. ^a^ PEAKS DB’s -10lgP threshold score was set as a cutoff filter to achieve false discovery rate (FDR) of 1.0% at the peptide sequence level.

| Species | Data set identifier | Parent Mass Error Tol. | Fragment Mass Error Tol. | Missed cleavages | Fixed modifications | Variable modifications | -10lgP threshold<sup>a</sup> | Instrument |
| --- | --- | --- | --- | --- | --- | --- | --- | --- |
| *Arabidopsis thaliana* | PXD000033 | ±5.0 ppm | ±0.5 Da | three | Beta-methylthiolation (+45.99 Da), iTRAQ 4plex (K, N-term) (+144.10 Da) | Oxidation (M) (+15.99 Da), Phosphorylation (STY) (+79.97 Da), Acetylation (N-term) (+42.01 Da), Deamidation (NQ) (+0.98 Da), iTRAQ 4plex (Y) (+144.10 Da) | 30 | Orbitrap (Orbi-Trap) |
|  | PXD000421 | ±7.0 ppm | ±0.8 Da | two | Carbamidomethylation (+57.02 Da) | Oxidation (M) (+15.99 Da), Phosphorylation (STY) (+79.97 Da), Acetylation (N-term) (+42.01 Da), Deamidation (NQ) (+0.98 Da) | 24.2 | Orbitrap (Orbi-Trap) |
| *Oryza sativa* Indica | PXD002222 | ±20.0 ppm | ±0.05 Da | one | Carbamidomethylation (+57.02 Da) | Oxidation (M) (+15.99 Da), Phosphorylation (STY) (+79.97 Da), Acetylation (N-term) (+42.01 Da) | 19 | Orbitrap (Orbi-Trap) |
|  | PXD000923 | ±15 ppm | ±0.05 Da | one | Carbamidomethylation (+57.02 Da) | Oxidation (M) (+15.99 Da), Phosphorylation (STY) (+79.97 Da) | 21 | Triple TOF |

### Extraction of Confidently Identified Phosphopeptides and Phosphoproteins

Code was written in Python (version 3.5.2) to merge exported PEAKS results (protein-peptides.csv file) for each species and extract confidently identified phosphopeptides and phosphoproteins. For the purposes of downstream analysis of sites of phosphorylation (phospho-sites), we merged peptide-spectrum matches (PSMs) identifying the same peptide and to ensure confident site analysis, we selected phosphorylation sites with Ascore ≥ 20. An Ascore of 20 will localize the site of modification with 99% certainty (p = 0.01) (34).

### Statistical analysis of the extent of multi-phosphorylation

Two-proportion z-tests were performed in R version 3.3.3 to test whether the observed proportion of multi-phosphorylated peptides in *Arabidopsis* is greater than the observed proportion in rice.

### Phospho-site Distance analysis

Proteins with more than two phospho-sites identified in the source data sets were extracted from our list of confidently identified phosphoproteins. Distances between phospho-sites were measured using a sliding window of 30 amino acids, starting from the first phosphorylated residue from the N-terminus. The distance was calculated by counting the number of amino acids until the next adjacent phosphorylated residue towards the C-terminus. Each pair of sites was assigned to a group based on the type of amino acid and the direction from the N-to-C termini. A background distribution, used to determine whether distances between phosphoserines observed differed from the random expectation, was calculated by two methods: one in which we calculated the distance between all serine residues in the theoretical proteome (all proteins in the species’ FASTA file), and one in which a peptide digestion model was included, to model the random sampling of peptides that occurs in LC-MS/MS workflows. However, both models produced near identical distributions (data not shown), and thus the former is presented for simplicity in interpretation.

### Phosphorylation Motif Prediction

Phosphopeptide sequences were submitted to Motif-x (35, 36) to predict phosphorylation motifs present in our identified phosphoproteins. 15-mers were constructed using Python, the pre-aligned 15-mer peptides were centered on each phosphorylated serine, threonine, and tyrosine and extended seven residues towards the N-terminus and seven residues towards the C-terminus. When the site was located less than seven residues from the N/C-terminus of the protein, the 15-mer was completed with the letter “X” to reach the required length of fifteen residues. For onward analysis, the occurrence threshold (i.e. number of 15-mer peptides with each motif) was >= 5, and the significance threshold <= 0.00018, to ensure a p-value of at least 0. 05 by the Bonferroni global correction method. *Oryza sativa* Indica and *Arabidopsis thaliana* proteomes obtained from UniProt version 59 were used to supply the background distribution for rice and *Arabidopsis* analyses, respectively. Motif-x results for *Arabidopsis* and rice were analyzed using Python code to identify motifs shared in the two species. A motif was classified as “shared”, if the same motif was identified passing the thresholds above in both species.

### Functional classification of motif containing proteins

Python code was written to obtain accessions for proteins from *Arabidopsis* and rice containing shared phosphorylation motifs. Protein accessions containing each shared motif were then grouped together into a list. The PANTHER statistical overrepresentation test (37) was used to classify each shared motif list into inferred biological processes, using the Bonferroni correction to apply global correction for multiple tests.

PANTHER uses *Oryza sativa* Japonica as a reference list for rice. To counteract the problem of using different cultivar, OrthoMCL (38) was first used to obtain Japonica accessions, then OrthoMCL results were analyzed (via Python code) to extract accessions orthologous to rice Indica. Biological processes with fewer than five mapped proteins were excluded from further analyses. PANTHER results were analyzed (via Python code) to extract biological processes that were inferred in both species by PANTHER for each shared motif list.

### Shared motif pair analysis

We next examined the “shared” phosphorylation motifs in both species independently, to examine whether particular pairs of motifs co-occurred within the same protein more often than would be expected by chance. All possible pair combinations of the twenty-one motifs were considered. Observed counts of co-occurrence less than five proteins were excluded from statistical analysis. The enrichment factor was calculated as the ratio of observed count of proteins that contains a shared motif pair with expected count across confidentially identified phosphoproteins. A one-tailed Fisher’s exact test was used to compare if the observed counts of protein containing a motif pair were significantly higher than expected values, using the 2×2 contingency table as shown in Table 2. The p-values for multiple testing were corrected by the Benjamini-Hochberg procedure.

**Table 2.** 2×2 contingency table was set to determine whether a pair of motifs occurred more often together than would be expected by chance where: q is the count of proteins with motifs a and b; m is the count of proteins with motif a; n is the count of proteins with motif b; N is the count of all identified phosphorylated proteins. Enrichment Factor (EF) calculated as EF_a,b_=q/(mn/N).

|  | Observed count | Expected count |
| --- | --- | --- |
| **Proteins with motif a and b** | q | $mn/N$ |
| **Proteins without motif a and b** | $N-m-n+q$ | $((N-m)(N-n))/N$ |

## Results and Discussion

### Identification of Phosphorylation Sites, Peptides, and Proteins

By combining the results from two datasets for each species and only selecting phosphopeptides with confidently identified phosphorylation sites, a total of 6,537 unique phosphopeptides from 3,189 unique phosphoproteins were identified in *Arabidopsis* (∼50% of the total proteins we identified). In rice, we identified 2,307 unique phosphopeptides from 1,613 unique phosphoproteins (56% of the total identified proteins in rice) – see Supplementary File 1 for all identifications.

To explore whether there are apparent global differences in the extent of phosphorylation across the two species, we profiled the numbers of phospho-sites per peptide and per protein. We identified a significantly greater proportion of singly phosphorylated peptides in rice (89.7%) than in *Arabidopsis* (78.9%) (Table 3). The majority of multi-phosphorylated peptides have two phosphates in both species. Only 2.3% of the phosphopeptides in *Arabidopsis* and 1.1% of the phosphopeptides in rice were suggested to have three phosphates. We found no phosphopeptides with four or more phosphates in this study, although this may be limited by overall the length of peptides identified.

**Table 3.**
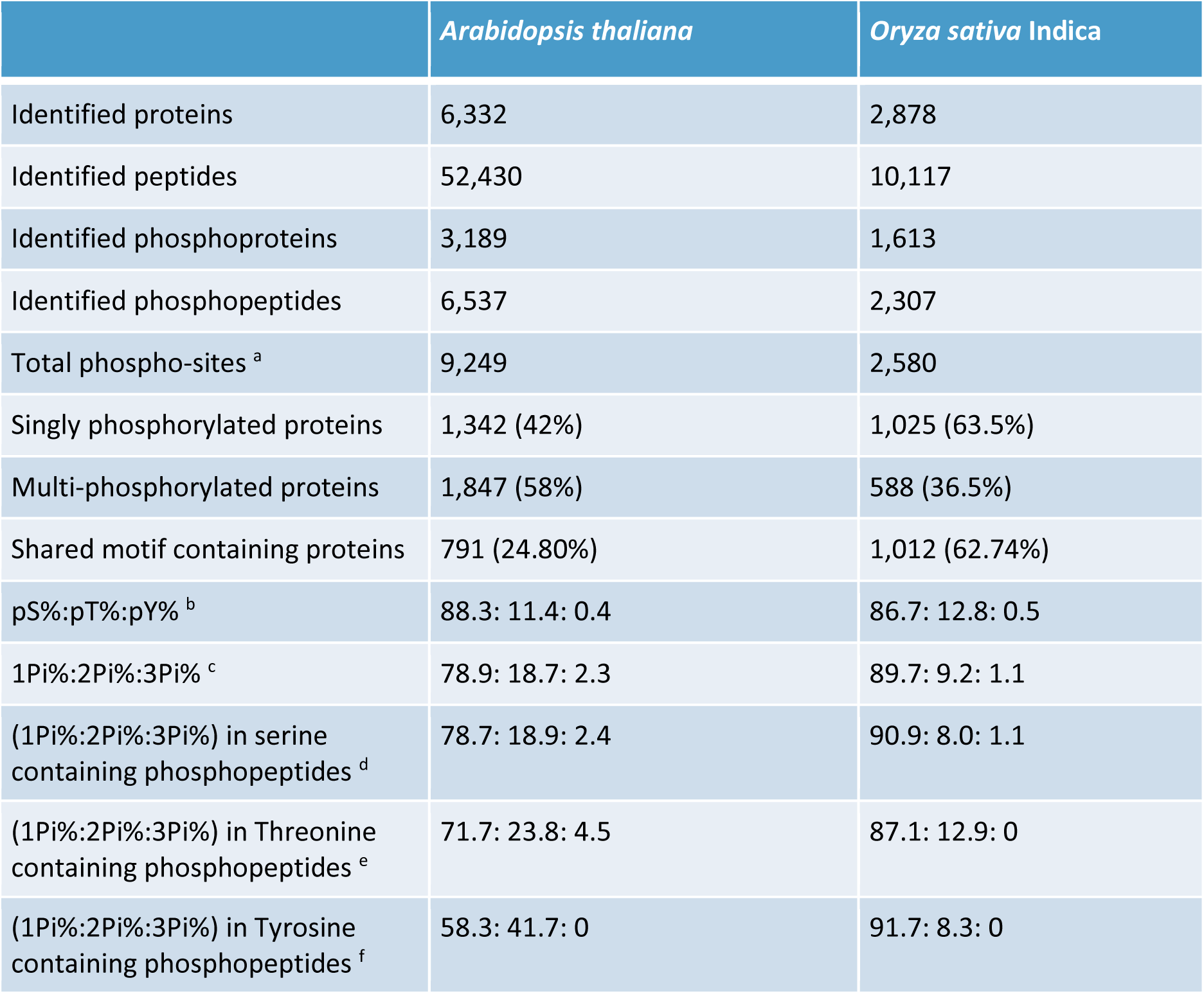
Summary statistics for protein and peptide identifications in our study. ^a^ Phosphorylation sites assignment with 99% certainty based on a p-value of 0.01 (Ascore ≥ 20); ^b^ Relative abundance of serine, threonine, and tyrosine phosphorylation sites based on analyzing 9,249 phosphorylation sites in *Arabidopsis* and 2,580 phosphorylation sites in rice.; ^c^ Relative abundance of single, double, and triple phosphopeptides based on analyzing a total of 6,537 p-peptides Arabidopsis and 2,307 p-peptides in rice - no phosphopeptides with four or more phosphates were found in this study. ^d^ Relative abundance of single, double, and triple phosphopeptides in serine containing phosphopeptides based on analyzing a total of 7,489 serine containing phosphopeptides in *Arabidopsis* and 2,148 serine containing phosphopeptides in rice. ^e^ Relative abundance of singly, doubly and triply phosphorylated peptides in threonine-containing phosphopeptides based on analyzing a total of 1,022 threonine containing phosphopeptides in Arabidopsis and 325 threonine-containing phosphopeptides in rice. ^f^ Relative abundance of singly, doubly and triply phosphorylated peptides in tyrosine containing phosphopeptides based on analyzing a total of 36 tyrosine containing phosphopeptides in *Arabidopsis* and 12 tyrosine containing phosphopeptides in rice.

Our results showed that the percentage of multi-phosphorylated peptides is significantly greater (by two fold) in *Arabidopsis* (21% i.e. summing values in 2Pi% and 3Pi% in Table 3) compared to rice (10.3%) with a p-value = 2.2e-16. The likelihood of observing multi-phosphorylated peptides has the potential to be influenced by experimental factors, including differences in the digestion of proteins into peptides (39) or the enrichment protocol (40). If proteins undergo less complete tryptic digestion, we would observe longer peptides, and thus the potential for more multi-phosphorylated peptides. To investigate whether peptide length is biasing the number of multi-phosphorylated peptides, we analyzed the length distribution for phosphorylated, non-phosphorylated and all peptides for rice and *Arabidopsis* (box plots for the distributions are presented in Supplementary Figure 1). The analysis shows that there is no overall difference between all peptides (rice versus *Arabidopsis*) – median length 14 amino acids in both cases (quartiles 11 - 19 amino acids). Phosphopeptides from rice (median: 13 amino acids; quartiles: 10 - 17) are slightly longer than *Arabidopsis* (median: 12; quartiles: 10 - 17), suggesting that peptide length does not explain why significantly more multi-phosphorylated peptides are observed in *Arabidopsis*. To explore whether differences in the enrichment protocols have caused more multi-phosphorylated peptides to be identified in *Arabidopsis* is not straightforward. However, one might hypothesize that a stronger enrichment for phosphopeptides over unmodified peptides might give rise to a higher proportion of multi-phosphorylated peptides. In *Arabidopsis*, 12% (6,537/52,430) of the total peptides we identified were phosphorylated compared with 23% in rice (2,307/10,117). As such, it does not appear that there is a stronger enrichment for phosphopeptides in *Arabidopsis*, which could explain why multi-phosphorylated peptides appear more common. However, we acknowledge that a comparison using matched protocols would be more ideal to rule out all possible sources of experimental bias. In summary, the data are suggestive of a difference in the rate of multi-phosphorylation between the two species, and thus the extent of crosstalk between nearby phospho-sites.

Sugiyama *et al*. reported that phospho-tyrosine containing peptides in *Arabidopsis* are frequently (75%) multi-phosphorylated (41). Our results showed that 41.7% of all observed pY-containing peptides are multi-phosphorylated, whereas 28.3% of pT-containing peptides and 21.3% of pS-containing peptides are multi-phosphorylated, as shown in Table 3. Whilst the extent of multi-phosphorylation on pY containing peptides is not as high as previously reported, it appears evident that phosphorylation on tyrosine is coordinated with other nearby phosphorylation events as an additional regulatory mechanism.

When examining rice data, we do not see evidence for higher extent of co-modification associated with tyrosine phosphorylation. The multi-phosphorylation percentages in serine, threonine, and tyrosine containing phosphopeptides are 9.1, 12.9 and 8.3% respectively. However, given the overall low counts of tyrosine containing phosphopeptides in rice, we cannot conclude that there is significant difference.

### Coordination in proximal phosphorylation sites in *Arabidopsis* and rice

We examined the distribution of distances between phosphorylated residues in multi-phosphorylated proteins from the N-to-C terminal direction (Figure 1). We have also scaled the frequency of serine to serine residues in all proteins, to act as a background distribution (for comparison with phosphoserine pair distances). Most of the paired phospho-sites follow the “randomly expected” trend from the background distribution with the exception of “pSXpS” (distance = 1), “pSXXpS” (distance = 2) and “pSXXXpS” (distance = 3), where there is evidently a strong enrichment in the paired phosphoserine sites in both species. There is no evidence for enrichment of a pSpS motif (distance = 0). The data thus demonstrate that in both rice and *Arabidopsis* there is functional crosstalk between phosphoserine sites separated by 1, 2 or 3 amino acids.

**Figure 1.**
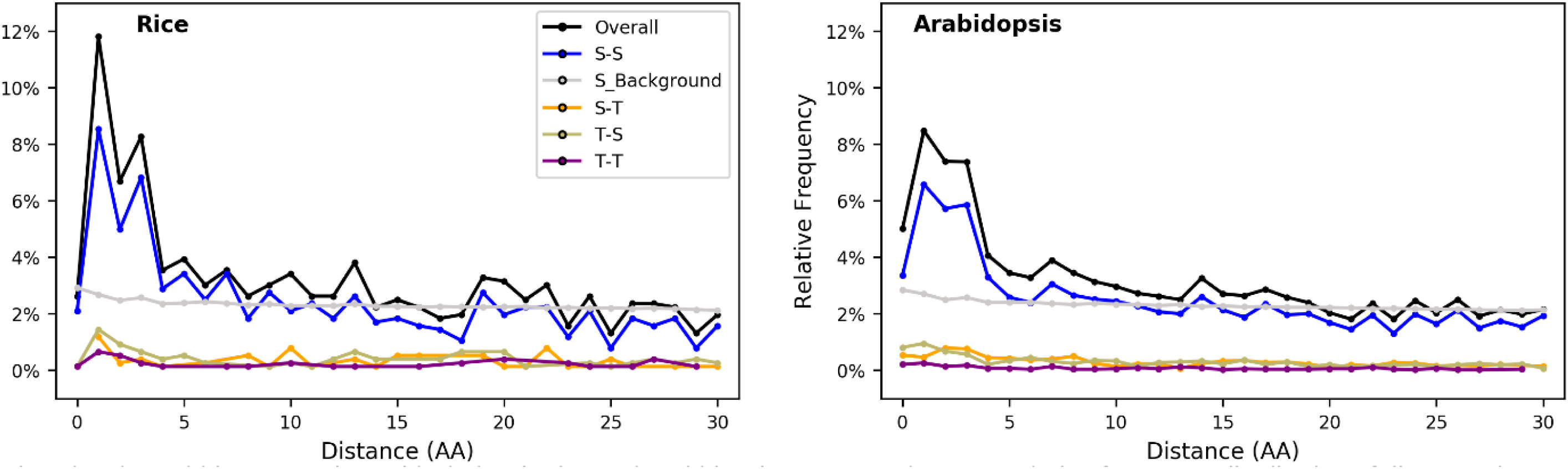
Relative frequency distribution of distances between two phospho-sites within a 30 amino acid window in rice (left) and *Arabidopsis* (right). In each subfigure, the gray line represents the background distributions in which we calculate as the distance between all serine residues in the theoretical proteome.

### Shared Motifs in *Arabidopsis* and Rice

We identified in total 76 pS, 7 pT, and one pY motifs in *Arabidopsis* and 51 pS, 6 pT with no pY motifs found in rice due to limited number of phosphotyrosines in this data set (Supplementary File 2). The most abundant pS motifs are RS in *Arabidopsis* and SP in rice (Figure 2). While TP, is by far the most common pT motif. Nineteen pS motifs and two pT motifs are shared in rice and *Arabidopsis* (making them candidate motifs likely to present in general in flowering plants) of which, four motifs (SP, TP, SxD, and SPR) are among the ten most abundant motifs predicted in rice and *Arabidopsis*, as shown in Figure 2. We grouped the pS motifs into types or families similar to van Wijk et al. (42), for example, TP-, SP-, SD-, GS-, T-, and S-types. Most of the pS shared motifs belong to the basic SP-type, four of which (RxxSP, SPK, SPR, SPxR) are found to be highly enriched in the *Arabidopsis* nucleus. While SPR found to be highly enriched in pistil tissue (female reproductive part of a flower) in rice (28). Motifs from the SD- and SP-types (SDxE, SP, SD) are very commonly targeted by Calcium-Dependent Protein Kinases (CDPKs). These Ser/Thr protein kinases are only found in plants and green algae (43) and have an essential role in plant defense response (44).

**Figure 2.**
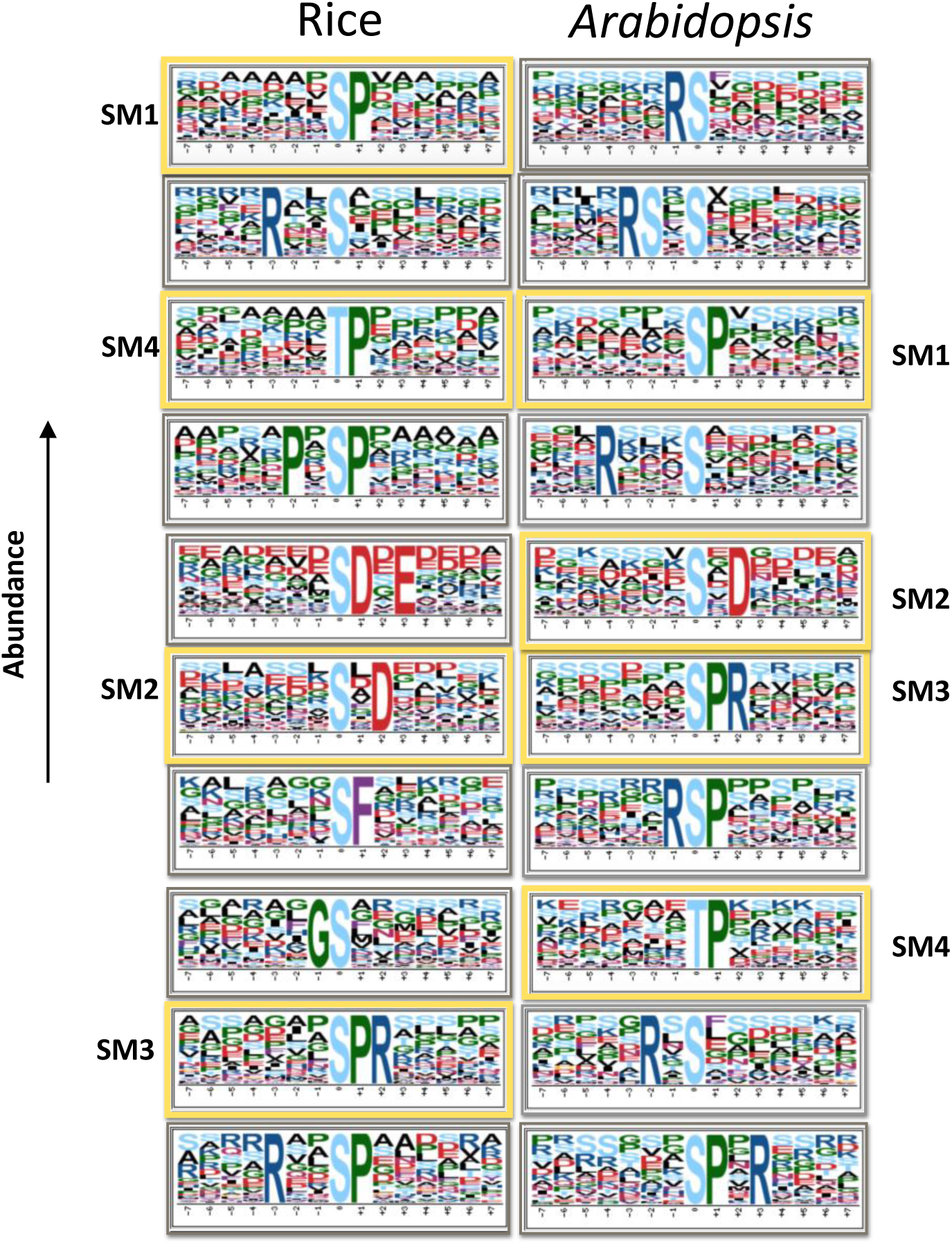
Motif logos of the ten most abundant motifs predicted in rice (left) and *Arabidopsis* (right). The logo indicates statistically significant residues at the p < 0.05 level corresponding to the 0.00018 significance threshold that was specified in motif-x analysis and the frequency of other residues that failed to exceed the significance threshold, yet may be statistically overrepresented. Framed in yellow, four motifs are shared (SM1-SM4) between the two species. Note that GS and RSP are shared motifs but not in the ten most abundant motifs in *Arabidopsis* and rice, respectively.

### Unique and Novel Motifs

Fifty-one pS, five pT, and one pY motifs are unique to *Arabidopsis* (i.e. we did not identify them in our rice data). Unique motifs such as RS, RSxS, and RxxxS are among the ten most abundant motifs in *Arabidopsis*. In rice, 29 pS and 4 pT motifs are considered as unique. Unique rice motifs such as SF, LxRxxS together with shared motifs SP, TP, and RxxS have been linked to rice response to bacterial blight (27). Many of the identified *Arabidopsis* motifs in our study overlap with motifs found in a recent meta-analysis of phosphoproteome datasets in *Arabidopsis* by van Wijk et al. (42) We also were able to identify RxxSF motif which was predicted by Wang et al. (45) but was not apparent in van Wijk *et al*. Fifty-five of the identified *Arabidopsis* motifs and forty-nine of the identified rice motifs in our study are novel motifs i.e. have not previously been identified in another study to our knowledge. Among them, three are shared motifs: KxxxSP, KxxxxSP, LxRQxS, see Supplementary File 2 for a full list of shared, unique, and novel motifs.

### Gene Ontology Classification by Biological Process

The majority of shared motif containing proteins are mapped to shared biological processes in rice and *Arabidopsis*, as shown in Figure 3. Similar patterns of fold enrichment for similar biological processes in rice and *Arabidopsis* were observed by each class of phosphorylation motif. Fold enrichment provides information of relatively how many more (or fewer) proteins in our test list map to a particular biological process than what would be expected by chance (46).

**Figure 3.**
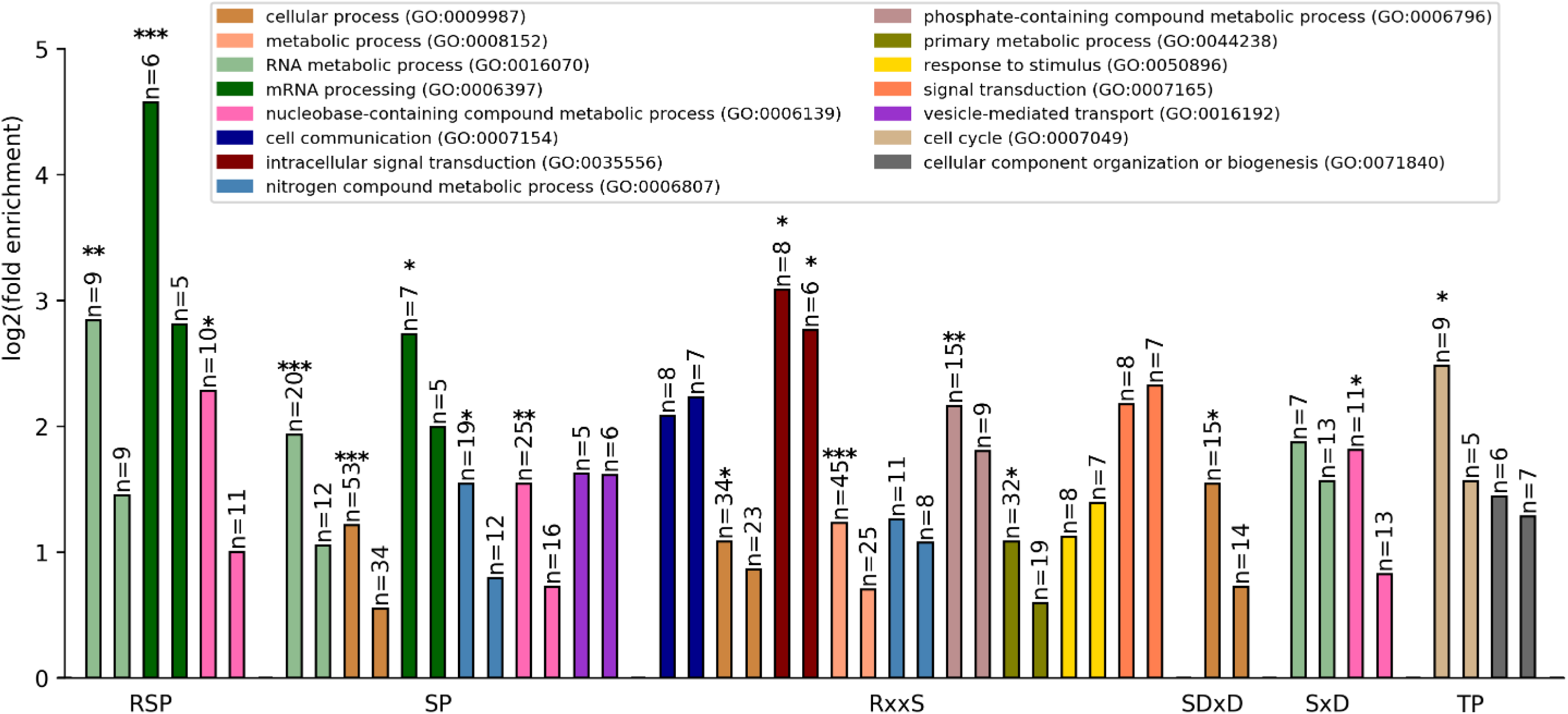
Log2(fold enrichment) for PANTHER GO-slim categories biological process terms mapped from rice (left bar of each pair) and *Arabidopsis* (right bar of each pair) phosphoproteins containing shared phosphorylation motifs. *= p< 0.05; **= p < 0.001; ***= p < 0.0001.

Proteins containing motifs of type RSP mapped in both species were enriched (statistically significant for rice only) for three related process category terms: “mRNA processing”, “RNA metabolism” and “nucleobase-containing compound metabolic process” - a high level process term including nucleobases, nucleosides, nucleotides and nucleic acid synthesis or metabolism. Proteins containing the related SP motif were mapped to the same processes, as well as “nitrogen compound metabolic process” (significantly enriched in rice only) – a term including proteins involved in nitrogen fixing, nitrification, denitrification and so on, and vesicle related transport (not significantly enriched in either species).

RxxS containing proteins were mapped to a wider set of biological processes including “intracellular signal transduction” and a variety of other fairly high-level terms relating to metabolic processes. Since the total numbers of proteins carrying each motif is comparatively low, high-level process categories have generally been extracted. It would only be possible to map to more specific process terms for very large counts of proteins. In general, the results indicate that shared motifs between rice and *Arabidopsis* map to the same biological processes, indicating that the association between kinase signaling and biological process are ancient, shared mechanisms in plants.

### Motif pair analysis

We examined the co-occurrence of shared motifs in *Arabidopsis* and rice (Figure 4). Proteins with at least two shared phosphorylated motifs were included in this analysis. Among shared motif-containing proteins (1012 in rice and 791 in *Arabidopsis*) more proteins with at least two shared phosphorylated motifs are found in *Arabidopsis* (340, 43%) compared to rice (268, 26.5%). Statistically significant enrichment was seen for several pairs of motifs in *Arabidopsis* including i) “RxxSP” with “SP”; ii) “RxxSP” with “SPxR”; iii) “SD” with “SxD”; iv) “KxxxxSP” with “SP”; v) “RxxSP” with “TP”; vi) “RxxSP” with “SPR”; vii) “SP” with “TP”; viii) “SDxD” with “SDxE”; ix) “GS with “RxxS”; x) “RSP” with “RxxSP”; xi) “SP” with “SPR”; xii) “SPK” with “TP”; xiii) “RSP” with “SPxR”; xiv) “SP” with “SxD”; xv) “KxxxSP” with “SP”; xvi) “SxD” with “TP”; xvii) “GS” with “SxD”; xviii) “GS” with “SP”; xix) “SP” with “SPxR”. It should be noted that in each case different sites have contributed to the motifs e.g. for cases where a motif is a subset of another “RxxSP” with “SP” the same sites did not contribute to both motifs. There is a trend towards motifs of the same general class e.g. SP type; SD or SxD to be most strongly enriched together. We interpret this to mean that the same kinase is likely phosphorylating multiple sites in the same protein, with similar motif types, rather than two kinases working in a coordinated manner. For cases where motifs of different classes are found to be co-occurring e.g. SP with SxD and GS with SxD, would point to protein sets likely under coordinated control from more than one kinase. As discussed above, we observe considerably fewer sites of multi-phosphorylation in rice than *Arabidopsis*, and this result is further exemplified in Figure 4. There are no motifs that co-occur in rice that pass statistical significance. It has previously been demonstrated in the human kinome that those proteins phosphorylated by a single kinase on a single site, participated in fewer pathways than those phosphorylated by multiple kinases (47). It is thus reasonable to assume that the expansion in both overall gene/protein count, and in the kinase count, in rice compared to *Arabidopsis*, leads to less crosstalk between phospho-sites in rice, and overall higher specificity in signaling through kinases. Further work is needed to explore the phenomenon across other plants species, so that research groups using *Arabidopsis* as a general model for plant biology and signaling can understand the extent to which findings are transferable.

**Figure 4.**
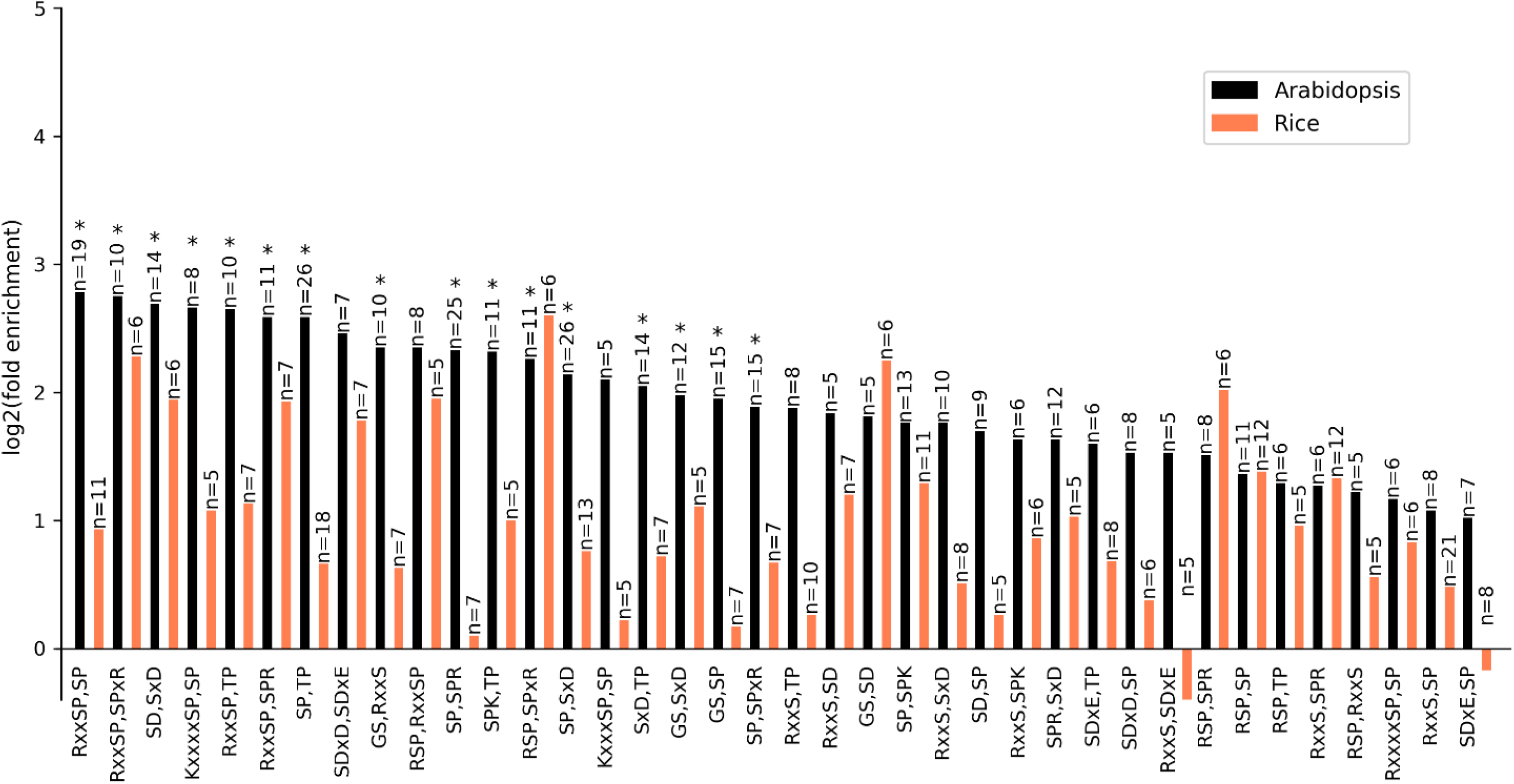
Co-occurrence of shared motifs in Arabidopsis and rice. The ratio of the observed-to-expected count of proteins that contain a motif pair from the twenty-one shared motifs identified in this study was calculated across confidentially identified phosphoproteins. n is the number of observed proteins for each shared motif pair.

## Conclusion

This study provided a thorough comparative analysis of the *Arabidopsis* and rice phosphoproteomes. By employing several datasets from previous phosphoproteomics studies, we confidently identified 6,537 phosphopeptides from 3,189 phosphoproteins in *Arabidopsis*, and 2,307 phosphopeptides from 1,613 phosphoproteins in rice, with the site of phosphorylation localized with 99% certainty. We identified 21 phosphorylation motifs shared between rice and *Arabidopsis*, and demonstrated further that they are associated with the same biological processes. It is reasonable to assume that phosphorylation-related functions are generally well conserved across diverse species of flowering plants, and that studies on the *Arabidopsis* as a key model species, can be used to make inferences about economically important crops, such as rice.

For both species, there is an enrichment for phospho-serine multi-phosphorylation sites separated by one, two or three amino acids. However, there are some differences between the two species (separated by ∼150 million years) in terms of overall gene and kinase count. We demonstrated that multi-phosphorylation was observed to be statistically higher in the *Arabidopsis* data than in the rice data, and we cannot found a confounding experimental factor that would explain the difference. Given the higher gene and kinase count in rice, it appears plausible that the difference is explained by higher specificity in signaling mechanisms in rice. Future studies on a wider range of model plant species should explore whether the functional implications of higher specificity and the extent to which this phenomenon extends to other plant species.

## Abbreviations

GO: Gene Ontology
pProtein: phosphorylated protein
pS: phospho-serine
pT: phospho-threonine
pY: phospho-tyrosine

## Acknowledgements

We are pleased to acknowledge funding from BBSRC and Newton fund that supported this work: [BB/N013743/1, BB/L005239/1].

